# Zika virus E protein alters blood-brain barrier by modulating brain microvascular endothelial cell and astrocyte functions

**DOI:** 10.1101/2023.02.09.527854

**Authors:** Guneet Kaur, Pallavi Pant, Reshma Bhagat, Pankaj Seth

## Abstract

Neurotropic viruses can cross the otherwise dynamically regulated blood-brain barrier (BBB) and affect the brain cells. Zika virus (ZIKV) is an enveloped neurotropic *Flavivirus* known to cause severe neurological complications, such as encephalitis and foetal microcephaly. In the present study, we used human brain microvascular endothelial cells (hBMECs) and human progenitor derived astrocytes to form a physiologically relevant BBB model. We used this model to investigate the effects of ZIKV envelope (E) protein on properties of cells comprising the BBB. E protein is the principal viral protein involved in interaction with host cell surface receptors, facilitating the viral entry. Our findings show that ZIKV E protein results in activation of both hBMECs and astrocytes. hBMECs showed reduced expression of endothelial junction proteins - ZO-1, Occludin and VE-Cadherin, which are crucial in establishing and maintaining the BBB. As a result, ZIKV E protein triggered alteration in BBB integrity and permeability. We also found upregulation of genes involved in leukocyte recruitment along with increased proinflammatory chemokines and cytokines upon exposure to E protein. Furthermore, E protein resulted in astrogliosis as seen by increased expression of GFAP and Vimentin. Both BBB cell types exhibited inflammatory response following exposure to E protein which may influence viral access into the central nervous system (CNS), resulting in infection of other CNS cells. Overall, our study provides valuable insights into the transient changes that occur at the site of BBB upon ZIKV infection.

## Introduction

Zika virus (ZIKV) is an enveloped, single stranded, positive sense RNA virus from the family *Flaviviridae*. It is an arthropod-borne virus spread through *Aedes aegypti* and *Aedes albopictus* mosquitoes and also transmitted through bodily fluids^1–3^. It is the only flavivirus known to cause teratogenic effects in humans, primarily resulting in abnormally small head *circumference*-microcephaly, intracranial calcification, and fetal death in some cases. ZIKV-associated clinical implications in adults predominantly include Guillain-Barré Syndrome (GBS), and other neurological complications such as encephalitis, meningitis, encephalopathy^4–7^. The genome of ZIKV is ~10.8 kb and encodes for single polyprotein which is later processed into three structural-capsid (C), envelope (E), premembrane/membrane (PrM), and seven non-structural proteins-NS1, NS2A, NS2B, NS3, NS4A, NS4B and NS5 ^8^.

The blood-brain barrier (BBB) is a complex and well-regulated interface between the peripheral and the central nervous system (CNS). BBB plays a pivotal role in maintaining the homeostasis of the CNS which includes - limiting passive diffusion of polar molecules from bloodstream into the brain, supply of nutrients and oxygen as well as efflux of harmful metabolites and xenobiotics, maintaining the water-electrolyte balance, and regulating the circulation of immune cells across the barrier^9,10^. BBB comprises of unique microvascular endothelial cells which line the cerebral capillaries associated with other cell types such as pericytes and astrocytic-end feet processes, which together form the ‘neurovascular unit’^11^. Brain microvascular endothelial cells (BMECs) are the primary component of BBB which are characterised by lack of fenestrations, scarce pinocytosis and presence of elaborate trans-membrane transport molecules^10–12^. Presence of tight junctions (occludin, claudin and zonula occludens-ZO) is the central feature of BMECs which are crucial in establishing and maintaining the BBB integrity. Inter-endothelial junctions also include adherens junctions (AJ), in which the primary component is vascular endothelium (VE)-cadherin. The mutual interactions between the cells comprising the neurovascular unit are crucial for barrier formation and maintaining its integrity.

A hallmark of CNS viral infections is disruption of BBB which can be induced by viral replication or results from neuroinflammation. Certain viral factors and host inflammatory responses can hamper BBB integrity. Other flaviviruses, Japanese Encephalitis virus (JEV) and West Nile virus (WNV) are known to disrupt BBB, whereas, the exact mechanisms involved in ZIKV neuroinvasion and encephalitis remain elusive. Disruption of BBB integrity in JEV and WNV infection resulted as cause of compromised tight junction complex^13,14^. ZIKV is detected in microcephalic brains of foetuses and is found to be neurotropic in adults with intact BBB^15^. The permissive nature of ZIKV towards cells comprising BBB needs further investigation.

Viral entry into host cells is facilitated by interaction of ZIKV envelope (E) protein with host surface receptors upon attachment. Evidences show the advent of ZIKV neuroinvasion and its virulence results from the genetic variation of 10 amino acids near the N-linked glycosylation site of E protein^16,17^. The N-linked glycosylation of E protein is a crucial determinant for its virulence. Asian ZIKV strains (H/PF/2013 and PRVABC59) are glycosylated at N154. This E glycosylation augments the transmission, viral attachment, and its pathogenesis^17,18^. In the current study, we employed ZIKV E protein of Asian strain (H/PF/2013) and for the first time we present experimental evidences as a consequence of virulent nature of ZIKV E protein with regard to its effects on the properties of human brain microvascular endothelial cells (BMECs) and human progenitor derived astrocytes (PDA), and its effect on BBB phenotype.

## Materials and Methods

### Human Primary Progenitor Derived Astrocytes

Human fetal brain tissue samples (10-14 weeks old) were collected with informed consent of the mother(s) from elective abortions. Samples were processed as per approved protocols by Institutional Human Ethics Committee in compliance with recommendations of Indian Council of Medical Research, New Delhi, India. Cells isolated from the telencephalon region were used to derive human neural progenitor cells (hNPCs) by culturing and passaging under aseptic conditions. hNPCs were then cultured on Poly-D-lysine (Sigma-Aldrich, St. Louis, MO, USA, Cat# P7280) coated culture flasks in neurobasal media (Invitrogen, San Diego, CA, USA, Cat# 21103-049) containing growth factors-epidermal growth factor (EGF, 20 ng/ml) (Peprotech, Rocky Hill, NJ, USA, Cat#AF-100-15-500UG) and fibroblast growth factor (FGF, 25 ng/ml) (Peprotech, Rocky Hill, NJ, USA, Cat#100-18B-50UG) supplemented with neuronal survival factor-1 (NSF-1) (Lonza, Charles City, IA, USA, Cat# CC-4323), N2 supplement (Invitrogen, San Diego, CA, USA, Cat# 17505-048), bovine serum albumin (BSA) (Sigma-Aldrich, St. Louis, MO, USA, Cat# A9418), glutamine (Sigma-Aldrich, St. Louis, MO, USA, Cat# G7513) and antibiotics-penicillin-streptomycin (Invitrogen, San Diego, CA, USA, Cat# 15140122), and gentamycin (Sigma-Aldrich, St. Louis, MO, USA, Cat# G1522). hNPCs were characterised by expression of their respective functional markers where more than 99% cells were positive for Nestin and SOX-2 (Supplementary figure **S1** B-G). These cells were further assessed for their ability to differentiate into neurons. More than 95% of differentiated cells were positive for neuronal markers MAP2 and Tuj1 (Supplementary figure **S1** H-M).

hNPCs were differentiated into an astrocytic-lineage by replacing with astrocyte growth media-complete minimal essential media (CMEM) (Sigma-Aldrich, St. Louis, MO, USA Cat# M0268-10X1L) supplemented with 10% fetal bovine serum (FBS) (Gibco, California, USA, Cat# 10270-106). The differentiation was continued for 21 days with half media change and splitting. More than 95% cells were immunopositive for Vimentin and glial fibrillary acidic protein (GFAP) (Supplementary figure **S1** N-S). The mature progenitor derived astrocytes (PDA) were used for further experiments. At least three different fetal tissues were used for the study.

### Human Primary Brain Microvascular Endothelial Cells

Primary human cerebral cortex microvascular endothelial cells (Passage 3, 12 CPD *in vitro,* ACBRI 376) were procured from Cell Systems (Kirkland, WA 98034, USA). Brain microvascular endothelial cells (BMECs) were cultured in Complete Classic Media (Kirkland, WA 98034, USA Cat#4Z0-500) supplemented with culture boost (Kirkland, WA 98034, USA Cat#4CB-500) and antibiotics-penicillin-streptomycin and gentamycin. Cells were grown in Attachment Factor (Kirkland, WA 98034, USA Cat#4Z0-210) coated culture-ware. Media was changed every two days and cells were split (> 80% confluent) with Passage Reagent Group (Kirkland, WA 98034, USA Cat#4Z0-800). Cultured hBMECs were found to be immunopositive for characteristic VE-Cadherin and ZO-1 (Supplementary figure **S2** A-F). Cells were used at passage 8-12 for different assays.

### Establishment of contact-based human Blood Brain Barrier model system using co-culture of Primary Progenitor Derived Astrocytes and Primary Brain Microvascular Endothelial Cells

For BBB monoculture and/or co-culture, Polyester (PET) (3.0 μm pore, 12 mm diameter, collagen-coated, Corning Life Sciences, ME, USA Cat# 3462) Transwell inserts were used. For establishing co-culture, transwell inserts were inverted and a new external well was created using a small piece of sterile elastic silicon tubing. The luminal side was sealed with smaller enclosed silicon tubing to avoid leakage of media. 35,000 PDAs were seeded onto the abluminal side of inserts and were allowed to adhere for 5-6 hours in CMEM. The inserts were inverted back to its upright position after removing the tubing gently, and cells were allowed to grow for 24 hours in CMEM. Next day, 75,000 transfected (with empty vector or ZIKV E-protein) BMECs were seeded onto the luminal side and both cell types were allowed to grow in ECs media for 24 hours and were harvested for different assays.

### Immunocytochemistry

Experiments were performed in eight-well and/or four-well Permanox chamber slides. Cells were seeded at 20,000 cells/ well and 50,000 cells/well, respectively. Cells were fixed with 4% paraformaldehyde (PFA) for 20 minutes, followed by three washes with 1X PBS. Blocking and permeabilization was done with 4% BSA and 0.3% Triton-X-100. Cells were probed with respective primary antibodies overnight at 4°C: anti-ZO-1 (1:500, Invitrogen, San Diego, CA, USA, Cat# 61-7300), anti-VE-Cadherin (1:500, Abcam, Cambridge, UK, Cat# ab33168), anti-GFAP (1:2000, Dako, USA, Cat# Z0334), anti-GFP (1:2000, Abcam, Cambridge, UK, Cat# ab290) and anti-Vimentin (1:2000, Santa Cruz, USA, Cat# sc-6260). Cells were washed three times with 1X PBS and incubated with suitable fluorophore tagged secondary antibodies (1: 2000, Invitrogen, San Diego, CA, USA). Cells were washed with 1X PBS before mounting, using hard set mounting media containing DAPI (Vector Labs, Burlingame, CA, USA). 6-7 random images were taken using AxioImager.Z1 microscope (Zeiss, Germany) from each group by person blinded to the experimental groups.

### Transient expression of ZIKV E protein

Full length ZIKV E protein cloned in pCAGIG-IRES-GFP expression vector was a kind gift from Dr Shyamala Mani (IISc, Bangalore, India) also used in previous studies^19,20^. Cells (80% confluent) were transfected using lipofectamine 3000 (Invitrogen, San Diego, CA, USA, Cat# L3000008) according to the manufacturer’s protocol. Cells were harvested after 24 hours transfection for further experiments (Supplementary figure **S3**). Empty vector was used as control.

### Transendothelial Electrical Resistance (TEER) Assay

Endothelial resistance was measured to assess BBB integrity of BMECs in monoculture as well as in co-cultured PDA and BMECs on transwell inserts. TEER was measured using Millicell ERS-2 electrical resistance instrument (Millipore, USA). After 24 hours, co-culture was established and measurements were done. Final calculations were done by multiplying TEER values with the area of insert membrane (1.1 cm2). TEER values of E protein transfected BBB culture were compared with monoculture/co-culture transfected with empty vector.

### Transendothelial Permeability Assay

PDA and BMECs grown on PET transwell insets were used to assess endothelial permeability. Dextran-FITC (M.W. 3000-5000) (Sigma, USA, Cat# FD4) was added to the upper compartment of insert at a final concentration of 50 μg/ml. The inserts were incubated at 37°C in dark for 30 minutes. Samples were then removed from the lower compartment for measuring fluorescence intensity using microplate fluorometer (E_x_ 480 nm and E_m_ 530 nm, Tecan).

### Western blotting

Protein extract was isolated using SDS lysis buffer (50 mM Tris Buffer (pH 7.5), 150 mM sodium chloride, 50 mM sodium fluoride, 1mM EDTA (pH 8.0), 2% SDS, 10mM sodium borate, 1mM sodium orthovanadate, protease inhibitor tablets (Roche, Basel, Switzerland, Cat# 11836170001). Protein concentration was estimated using 4% copper sulfate and bicinchoninic acid (Sigma, USA, Cat#B9643). Protein samples were separated by 8-12% SDS-PAGE. Proteins were then transferred onto nitrocellulose membrane (MDI, India), followed by 2 hours blocking with 5% skimmed milk at room temperature (RT). The blots were incubated with respective primary antibodies: anti-ZIKV-E protein (1:2000, GeneTex, Cat# GTX133314), anti-ZO-1 (1:2000, Invitrogen, Cat# 61-7300), anti-VE-Cadherin (1:2000, Abcam, Cat# ab33168), anti-Occludin (1:500, Invitrogen, Cat# OC-3F10), anti-GFAP (1:60000, Dako, USA, Cat# Z0334), anti-Vimentin (1:10000, Santa Cruz Biotechnology, USA, Cat# sc-6260), and anti-β-actin (1:40000, Sigma Cat# A3854), overnight at 4°C. The blots were washed with 1X TBST thrice for 5 minutes and then incubated with HRP-conjugated secondary antibodies (1:4000, Vector Labs, Burlingame, CA, USA) for 1-2 hours at RT. The blots were then washed using 1X TBST five times for 5 minutes. Protein was detected using chemiluminescence reagent (Millipore, Bedford, MA, USA, Cat# WBKLS0500 and imaged using Nine Alliance mini-HD UVITEC (Cambridge, UK). Protein bands so obtained were densitometrically quantified using ImageJ software (NIH, USA).

### Quantitative real-time PCR

Total RNA was extracted from transfected samples using TRIZOL (Ambion, Texas, USA Cat# 15596018) according to the manufacturer’s protocol. cDNA was synthesized from RNA using high-capacity cDNA reverse transcription kit (Applied Biosystems, Austin, TX, USA, Cat# 4368814) as per the manufacturer’s instructions. RT-PCR was performed using SYBR Green master mix (Applied Biosystems, USA, Cat# 4367659) using specific primers for IL-6, IL-8, IL-1β, CCL2, CCL5, CXCL10, ICAM-1, VCAM-1, PTGS2, GFAP, Vimentin and GAPDH, as mentioned in the supplementary table 1. The cycling conditions used were 95°C for 10 min (1 cycle), 95°C for 20s, 58°C for 20s, and 72°C for 30s (40 cycles).

### Cytokine Bead Array

BMECs were transfected with ZIKV E-protein and respective empty vector (control) for 24 hours, followed by collection of supernatants. To determine cytokine and chemokine levels, supernatant was incubated with cytokine beads (CBA; Multiplex magnetic bead-based antibody detection kits, BD Biosciences, CA, USA, Cat# 551811) as per the manufacturer’s protocol. Incubated complex was passed by flow cytometer, and represented data was analysed using the BD FACSDiva software.

### Monocyte chemoattractant protein-1 (MCP-1)/CCL2 ELISA

Cells were transfected with ZIKV E protein and empty vector, as control, for 24 hours. Supernatant was collected to determine the levels of MCP-1/CCL2 using commercial kit (BD Biosciences, CA, USA, Cat# 555179) as per manufacturer’s protocol.

### Extracellular Glutamate release

Astrocytes were transfected with ZIKV E-protein and respective empty vector (control) for 24 hours, followed by collection of supernatants. It was further processed for the detection of glutamate release using glutamate determination kit (Sigma Aldrich, USA, Cat# GLN1) as per manufacturer’s protocol.

### Statistical Analysis

Experimental results are represented as mean values ± standard error of the mean. Each experiment was performed at least three times to determine the significance of the means. Comparison between experimental and control group was analysed using Student’s t-test. A level of p< 0.05 was considered statistically significant.

## Results

### ZIKV E protein results in modulation in BBB integrity and permeability

Human brain microvascular cells (hBMECs) have been reported to show productive ZIKV infection^21–23^. ZIKV infects and activates hBMECs, however the effects of its surface protein-envelope (E) are not explored. In the present study, we examined the effect of ZIKV E protein on human brain microvascular endothelial cells (hBMECs) and astrocytes, and their resulting effect on BBB integrity. hBMECs were subjected to 24 hours transfection with ZIKV E protein and compared with cells transfected with empty vector (Supplementary figure **S3**). To investigate the effect of ZIKV E -protein on the integrity of endothelial barrier, quantitative measurement of TEER was done in hBMECs monoculture (Fig1. A). E-protein resulted in significant effect on the integrity of BBB in monoculture of hBMECs, as shown by decreased TEER values (135.1 ± 2.725, p <0.01) when compared with empty vector control (Fig. 1B). We also assessed transendothelial permeability of BBB in monoculture towards dextran-FITC upon exposure to ZIKV E protein. Monoculture layer of BMECs showed increased permeability to dextran-FITC (1.102 ± 0.019, p < 0.05) upon exposure to ZIKV E protein as compared to its control (Fig. 1C) We further investigated the impact of astrocytes on the integrity of BBB in a contact-based co-culture (Fig1. D). Presence of astrocytes with hBMECs resulted in higher TEER values compared to monoculture, suggesting that contact-based BBB model gives greater stability Albeit the transient expression of ZIKV E protein in hBMECs, decrease in TEER values (156.9 ± 3.748, p < 0.01) were observed in co-culture of BBB (Fig. 1E). Co-culture model of BBB also exhibited increased permeability towards dextran-FITC (1.052 ± 0.007, p < 0.01) (Fig. 1F). Permeability changes in BBB were more prominent in monoculture, as presence of astrocytes in the co-culture BBB model reflected greater barrier strength. These findings indicate perturbed BBB integrity as seen by decreased TEER values and increased permeability to dextran-FITC in both hBMECs monoculture and co-culture of hBMECs and astrocytes in response to E protein.

**Figure 1:**
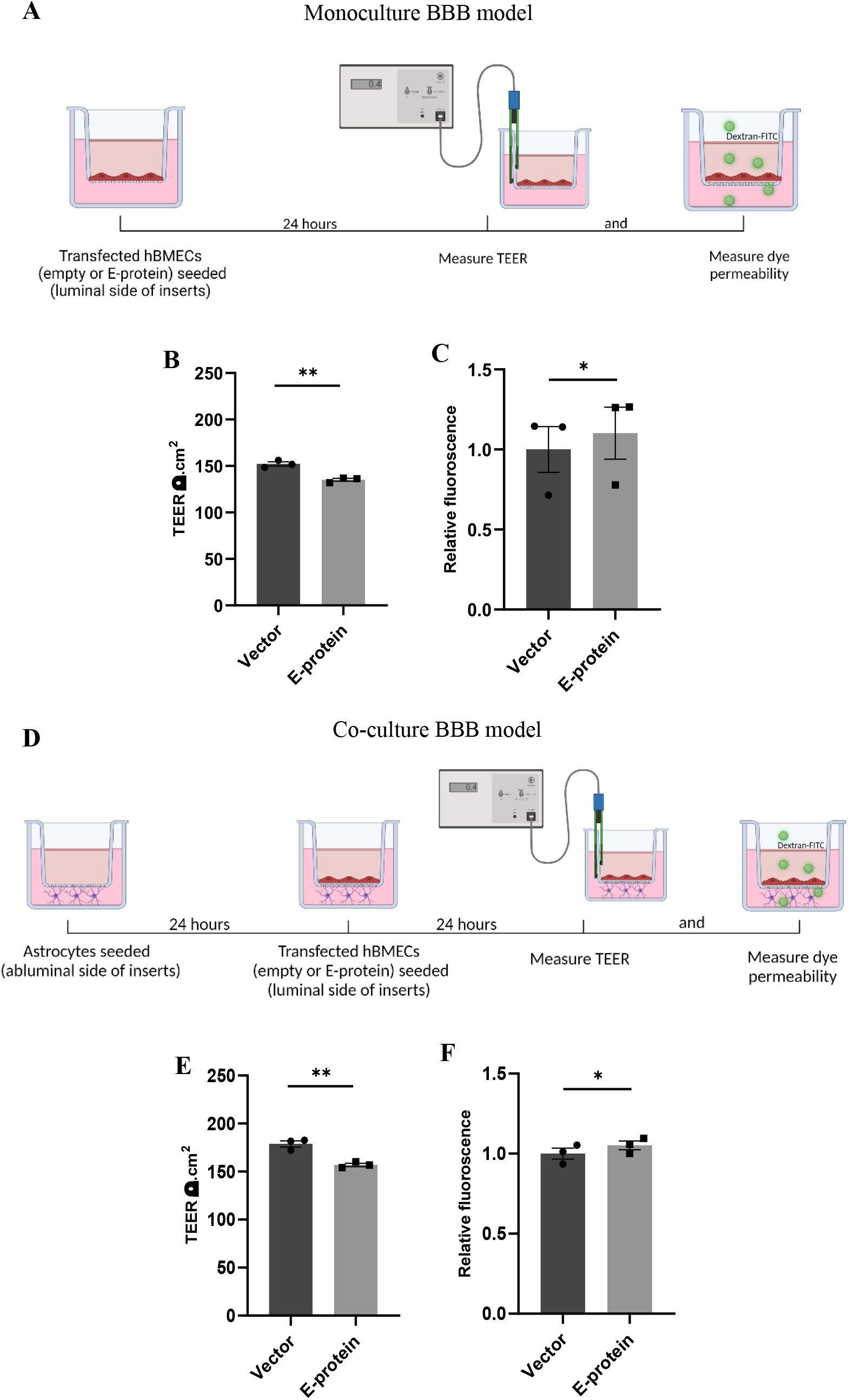
Effect of ZIKV E protein on BBB integrity. Schematic representation of monoculture BBB model establishment using hBMECs (A). hBMECs were transfected with empty vector or ZIKV E-protein. TEER was measured after 24 hours. Bar graph shows TEER values given in ohm.cm^2^ (B). Dextran-FITC transendothelial permeability was assessed after 24 hours of transfection. Bar graph shows relative fluorescence change (C). Schematic representation of coculture BBB model establishment using hBMECs and astrocytes (D). hBMECs transfected with empty vector or ZIKV E protein were grown in contact with astrocytes for 24 hours followed by measurement of TEER values (E) and dextran FITC transendothelial permeability (F). Data represents mean ± SEM for at least three independent experiments. *p<0.05, **p< 0.01 with respect to control.

### ZIKV E protein dysregulate expression of tight junction and adherens junction proteins in hBMECs

To investigate the molecular drivers of disrupted BBB induced by ZIKV E protein, we investigated the effect of ZIKV E protein on expression of tight junction and adherens junction proteins which are crucial in establishment and maintenance of BBB integrity. hBMECs were transfected using ZIKV E protein expression vectors for 24 hours and then harvested for further experimentation. We studied the expression of endothelial junction proteins, VE-cadherin, ZO-1 and Occludin by Western blotting. Western blotting results showed reduced levels of endothelial ZO-1 (0.724 ± 0.080, p < 0.05), VE-cadherin (0.630 ± 0.081, p < 0.01), and Occludin (0.806 ± 0.075, p < 0.05) proteins in ZIKV E-transfected hBMECs as compared to control cells which were transfected with empty vector (Fig. 2 A-D). These findings suggest that ZIKV E protein modulates the properties of hBMECs by altering expression of endothelial junction proteins.

**Figure 2:**
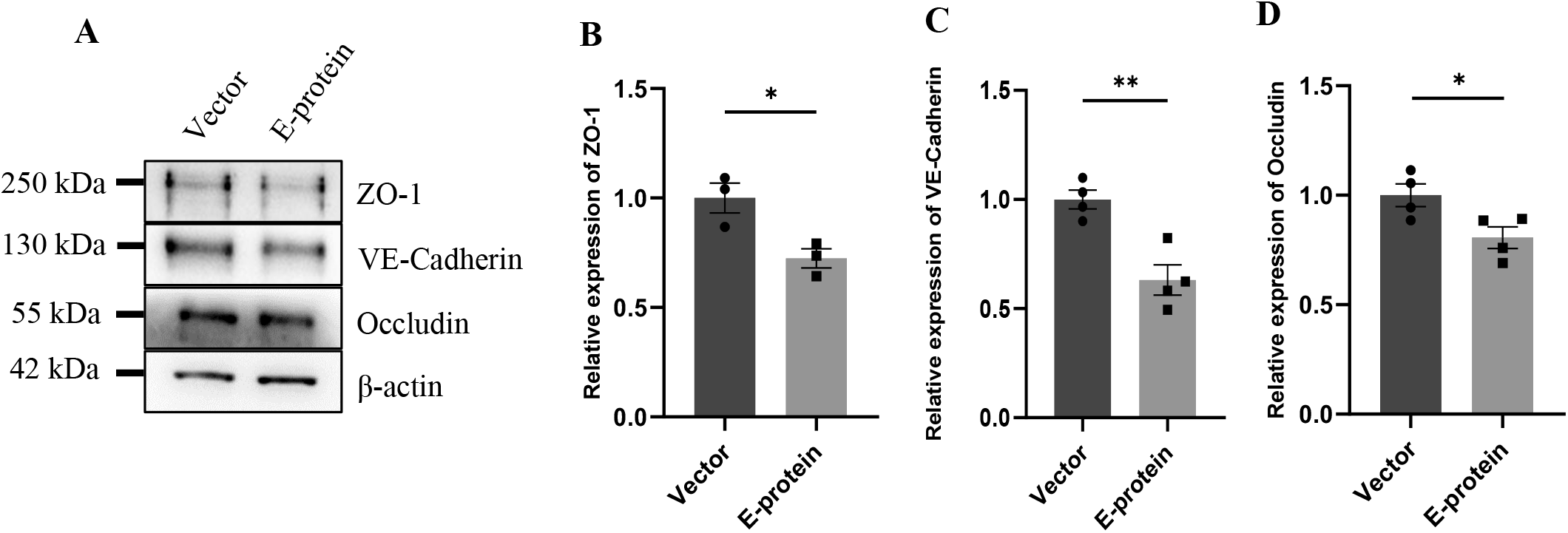
Effect of ZIKV E protein on endothelial junction protein expression. hBMECs were transfected with ZIKV E protein and empty vector as control for 24 hours. (A) Total protein was isolated and subjected to Western blotting against ZO-1 (B), VE-cadherin (C), occludin (D) antibodies. β-actin was used as loading control. One representative blot is shown. Bar graphs shows relative protein expression in indicated groups. Data represents mean ± SEM for at least three independent experiments. *p< 0.05, **p< 0.01 with respect to control.

### ZIKV E protein induces inflammation in hBMECs

BBB disruption enables inflammatory response at the injury site enabling the recruitment of immune cells, thereby, exacerbating the pro-inflammatory scenario and resulting in local inflammation^24,25^. We investigated effect of E protein on the endothelial homeostasis, by studying the expression of IL-6, IL-8, IL-1β, CCL2, CCL5, CXCL10, ICAM-1, VCAM-1 and PTGS2 genes in E protein and vector control transfected cells. Upregulation of genes involved in inflammation such as IL-6 (1.393 ± 0.098, p < 0.05), IL-8 (1.307 ± 0.083, p <0.05), IL-1β (1.868 ± 0.187, p < 0.01), CCL2 (1.937 ± 0.196, p <0.01), CCL5 (1.207 ± 0.079, p <0.05) and CXCL10 (1.504 ± 0.113, p < 0.01) was observed (Fig.3 A-F). Interestingly, genes involved in immune cell recruitment, ICAM-1(1.762 ± 0.254, p < 0.05), VCAM-1 (2.067 ± 0.197, p < 0.01) and angiogenesis, PTGS2 (1.720 ± 0.133, p < 0.001) were also significantly upregulated (Fig. 3 G-I). In addition to this, we examined the secretion of key proinflammatory cytokines IL-6 and IL-8 in the extracellular milieu of hBMECs transfected with E -protein. Increased secretion of IL-6 (1.417 ± 0.102, p < 0.01) and IL-8 (1.065 ± 0.019, p < 0.01) was observed as compared to control cells (Fig. 4 A-C). However, there was no difference in levels of IL-1β, IL-12p70 and TNF-a (data not shown). Our findings show perturbed endothelial homeostasis in response to E protein which can further modulate the expression and localization of tight junction and adherens junction proteins.

**Figure 3:**
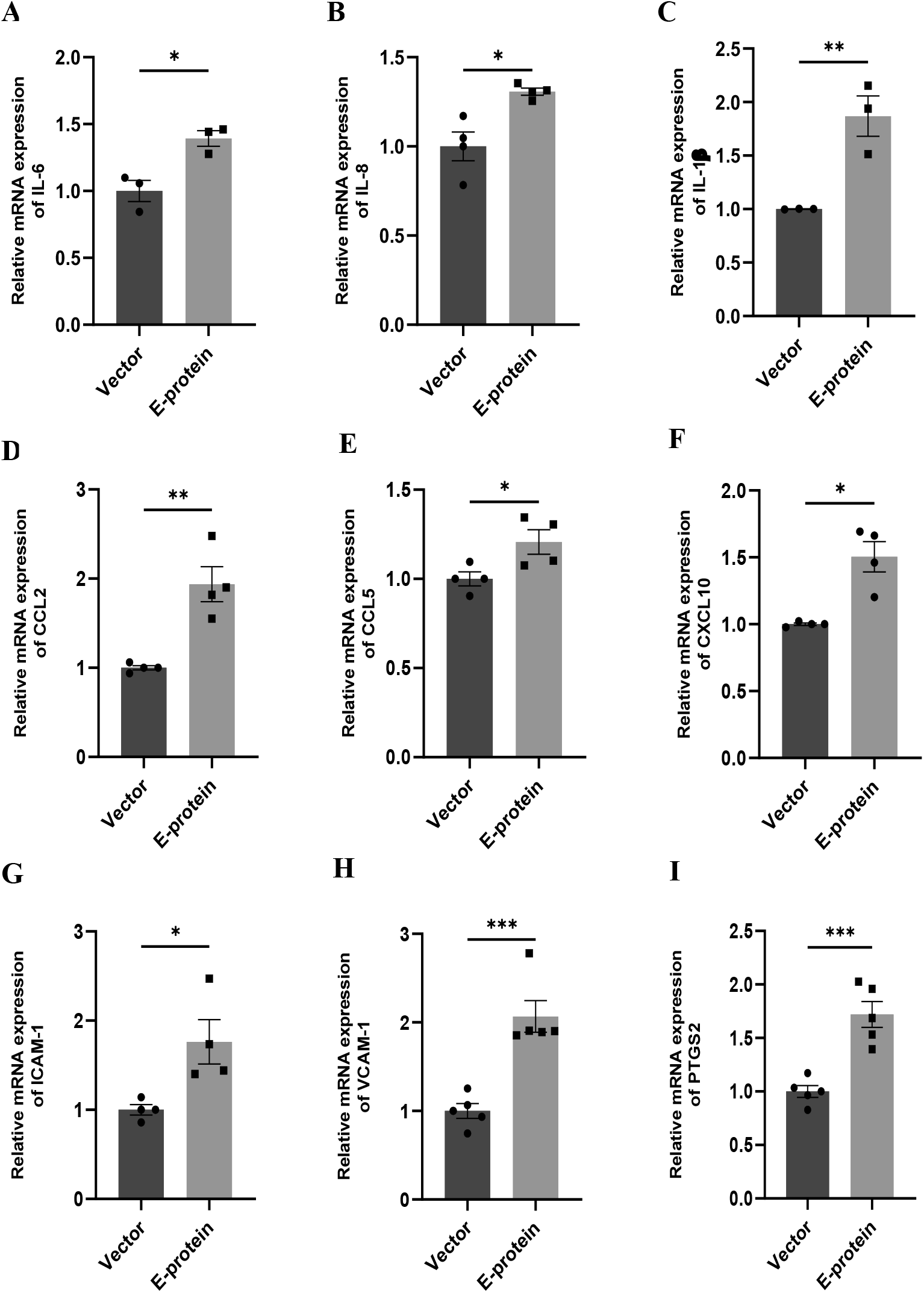
ZIKV E protein induces inflammation in hBMECs. Brain microvascular endothelial cells were transfected with empty vector and E protein for 24 hours. Total RNA was isolated and transcript levels of inflammatory cytokines and chemokines were measured using quantitative RT-PCR for IL-6 (A), IL-8 (B), IL-1β (C), CCL2 (D), CCL5 (E) and CXCL10 (F). Transcript levels of genes involved in modulation of adhesion molecules and angiogenesis-ICAM-1 (G), VCAM-1 (H), and PTGS2 (I) were also measured. Bar graph shows relative transcript expression in indicated groups. Data represents mean ± SEM for at least three independent experiments. *p<0.05, **p<0.01, ***p<0.001 with respect to control.

**Figure 4:**
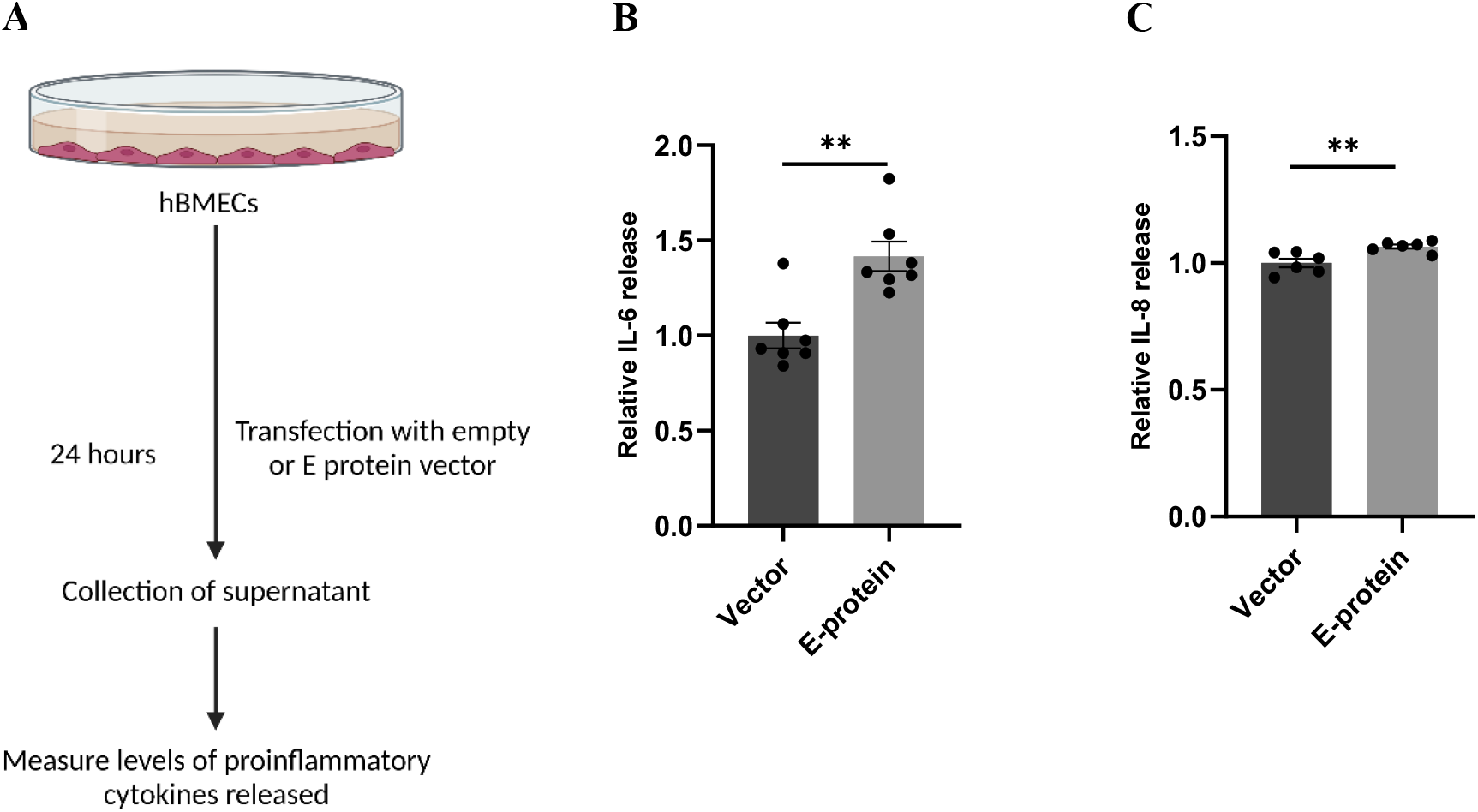
ZIKV E protein induces release of proinflammatory cytokines in hBMECs. Schematic representation of experimental setup (A). Brain microvascular endothelial cells were transfected with empty vector or E protein for 24 hours. Supernatants collected from transfected cells were subjected to cytokine bead array for measurement of inflammatory cytokines IL-6 (B) and IL-8 (C). Data represents mean ± SEM for at least three independent experiments. **p<0.01 with respect to control.

### ZIKV E protein triggers inflammatory response in astrocytes

Astrocytes are the major brain cell population and are known to be highly permissible towards ZIKV infection. Because of their involvement in BBB formation and its maintenance, we investigated the how ZIKV E protein impacts astrocytic activity. In response to CNS infections, astrocytes undergo morphological and functional changes pertaining to process known as “astrogliosis” or “reactive gliosis”^26^. Reactive astrocytes are characterized by increased levels of glial fibrillary acidic protein (GFAP) and vimentin^27^. Dysregulated release of various soluble factor by reactive astrocytes also corroborates towards neuropathologies^28^. We assessed the reactive state of astrocytes transfected with E protein and empty vector as control, for 24 hours. A remarkable upregulation of GFAP (2.252 ± 0.432, p < 0.05) and vimentin (1.990 ± 0.249, p < 0.05) was revealed in Western blotting (Fig. 5 A-D). Similar changes were also observed at the mRNA levels of GFAP (1.313 ± 0.135, p < 0.05) (Fig. 5E) and vimentin (1.919 ± 0.169, p < 0.01) (Fig. 5F).

**Figure 5:**
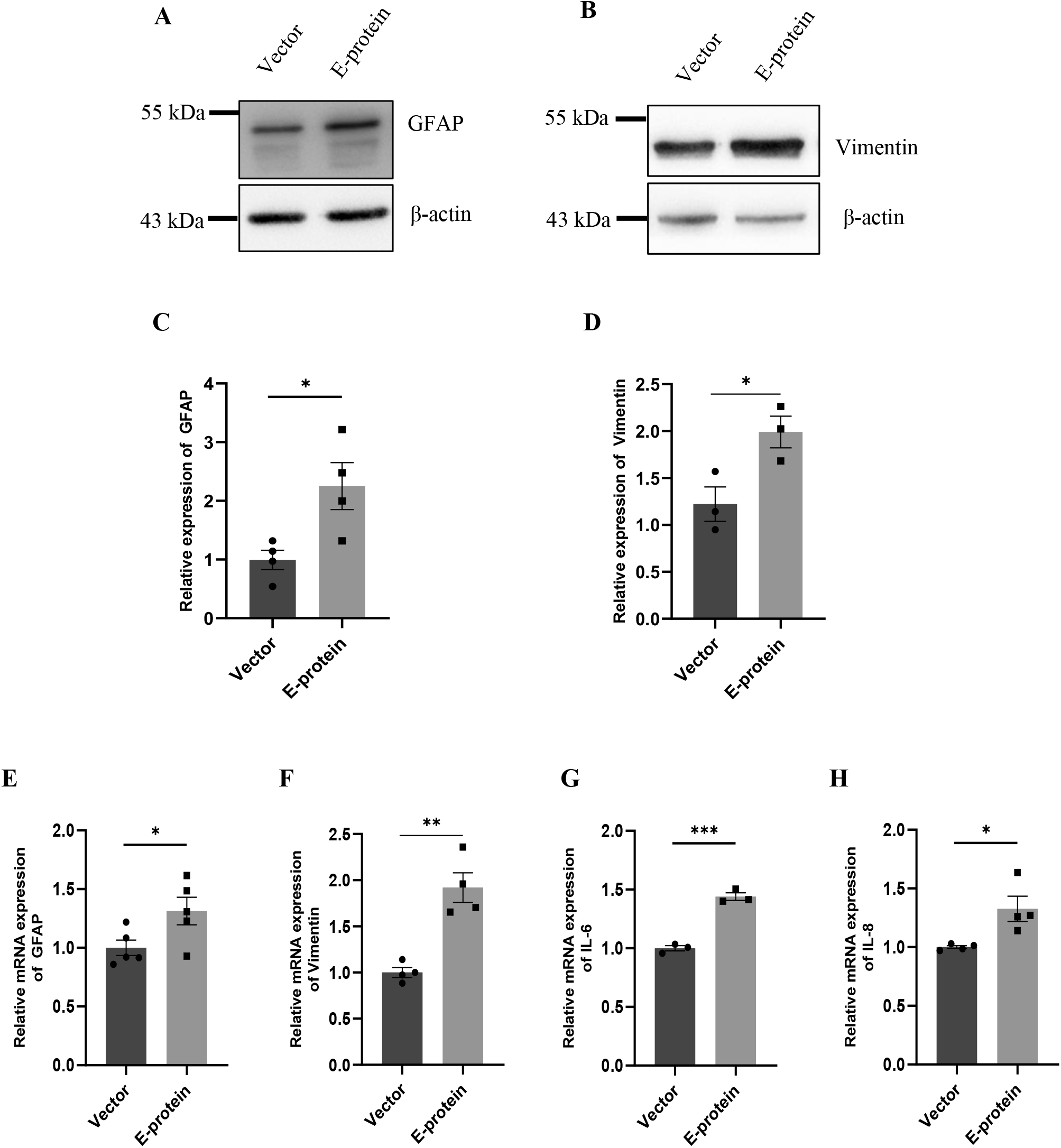
ZIKV E protein triggers astrogliosis. Progenitor derived astrocytes were transfected with empty vector or E protein for 24 hours. Total protein was isolated and subjected to Western blotting against markers for astrocyte reactivity, GFAP (A) and vimentin (B). β-actin was used as loading control. One representative blot is shown. Bar graph shows relative protein expression in indicated groups (C-D). To assess the transcript levels, total RNA was isolated and subjected to quantitative RT-PCR for GFAP (E), Vimentin (F), IL-6 (G), and IL-8 (H). Data represents mean ± SEM for at least three independent experiments. *p<0.05, **p<0.01, ***p<0.001 with respect to control.

We further investigated the inflammatory response of astrocytes towards ZIKV E protein expression. RT-qPCR showed elevated levels of IL-6 (1.439 ± 0.039, p < 0.001) and IL-8 (1.326 ± 0.108, p < 0.05) after 24h of ZIKV E protein expression (Fig. 5 G-H). These findings suggests that astrocytes exacerbated the inflammation at the BBB site, in response to ZIKV E protein. Furthermore, astrocytes actively contribute in maintaining glutamate homeostasis in the brain which is perturbed in the reactive state of astrocytes, thereby pertaining to glutamate excitotoxicity^29^. Hence, we wanted to study secretion of glutamate from astrocytes after the ZIKV E protein transfection. E protein transfected astrocytes resulted in increased levels of glutamate in the extracellular milieu (1.337 ± 0.059, p <0.01) (Fig. 6 A, B). Monocyte chemoattractant protein 1 (MCP-1) is one of the most commonly expressed chemokine expressed during CNS inflammation and is critical for monocyte recruitment and migration, BBB alteration, hence propagating inflammation^30^. We checked the levels of MCP-1 release in supernatant collected from transfected astrocytes. Results showed elevated levels of MCP-1 (2.002 ± 0.354, p < 0.05) (Fig. 6 A, C). These findings indicate astrocytes being essential site of ZIKV infection potentiates inflammation in its reactive state when exposed to ZIKV E protein. These results suggest that astrocytes play role in altering integrity of endothelial barrier.

**Figure 6:**
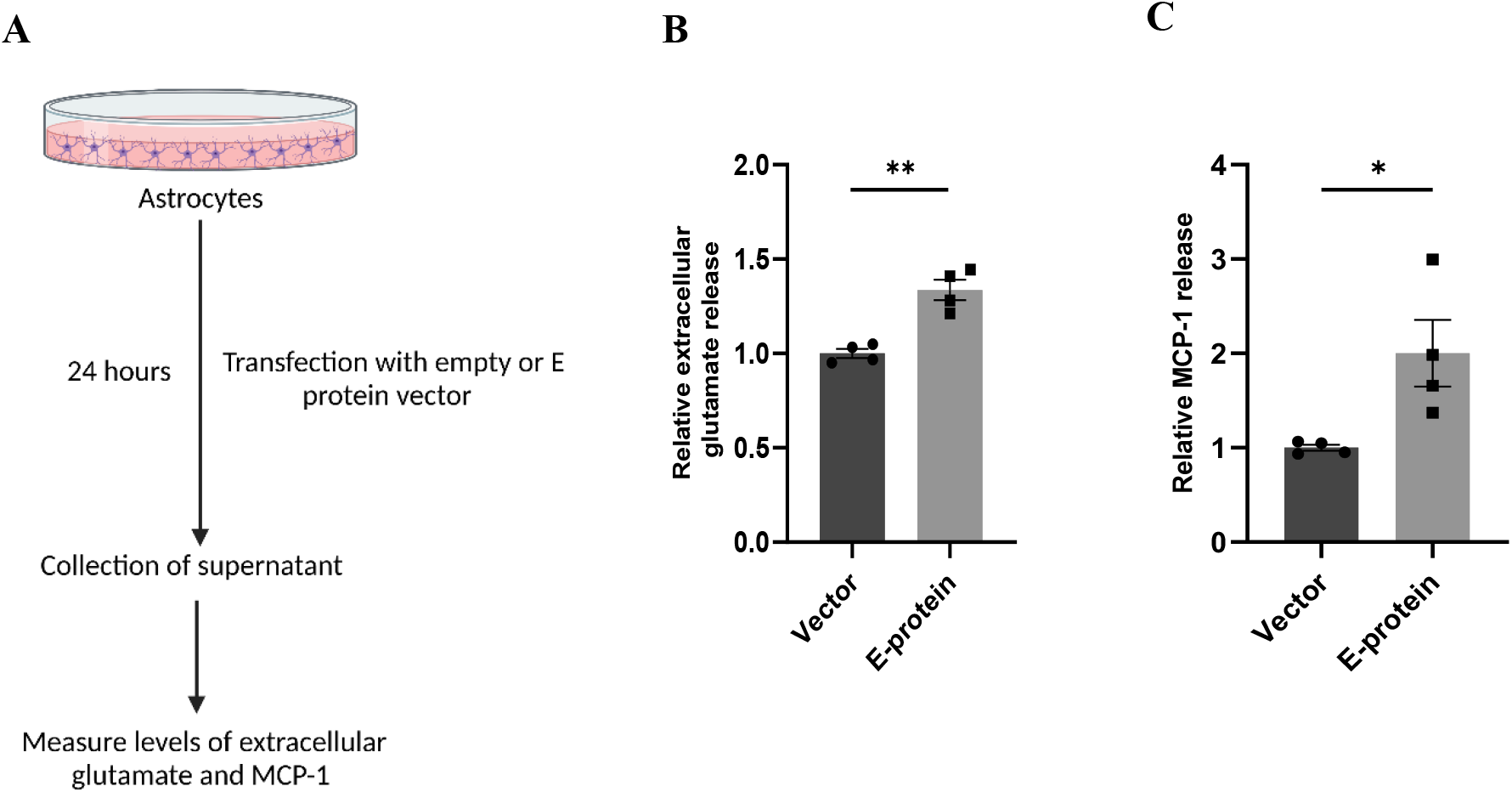
ZIKV E protein induced reactive astrocytes release extracellular glutamate and MCP-1. Progenitor derived astrocytes were transfected with empty vector or E protein for 24 hours. Schematic representation of experimental paradigm followed (A). Supernatant collected from transfected astrocytes was subjected to measurement of extracellular glutamate release (B) and also subjected to ELISA for measurement of MCP-1 (C). Data represents mean ± SEM for at least three independent experiments. *p<0.05, **p<0.01 with respect to control.

### ZIKV induced reactive astrocytes alter expression of endothelial proteins

As astrocytes and BMECs mutually interact with each other via soluble factors and enhance barrier properties^31^, we assessed whether E protein transfected ‘reactive astrocytes’ altered expression of endothelial junction proteins. To address this question, the supernatant from transfected astrocytes was collected after 24 hours and mixed with equal proportion of fresh endothelial media. hBMECs were then treated with astrocyte-conditioned media (ACM)-empty vector and/or E-protein, respectively.

E-protein exposure resulted in significant reduction in endothelial junction proteins-ZO-1 (0.641 ± 0.096, p < 0.01), VE-Cadherin (0.877 ± 0.196, p= 0.553), and Occludin (0.660 ± 0.075, p < 0.05) (Fig. 7 A-E). These results were further corroborated by our findings from contact-based co-culture model of BBB. Transfected hBMECs grown in direct contact with astrocytes were harvested after 24 hours to check expression of endothelial junction proteins. Results showed prominent reduction in ZO-1 (0.554 ± 0.099, p < 0.01), VE-Cadherin (0.563 ± 0.102, p < 0.05), and Occludin (0.413 ± 0.262, p < 0.05) (Fig. 8 A-E). These results strongly emphasize how mutual interaction of astrocytes and BMECs play important role in maintaining BBB integrity, as both cell types were found to be vulnerable to exposure to ZIKV E-protein.

**Figure 7:**
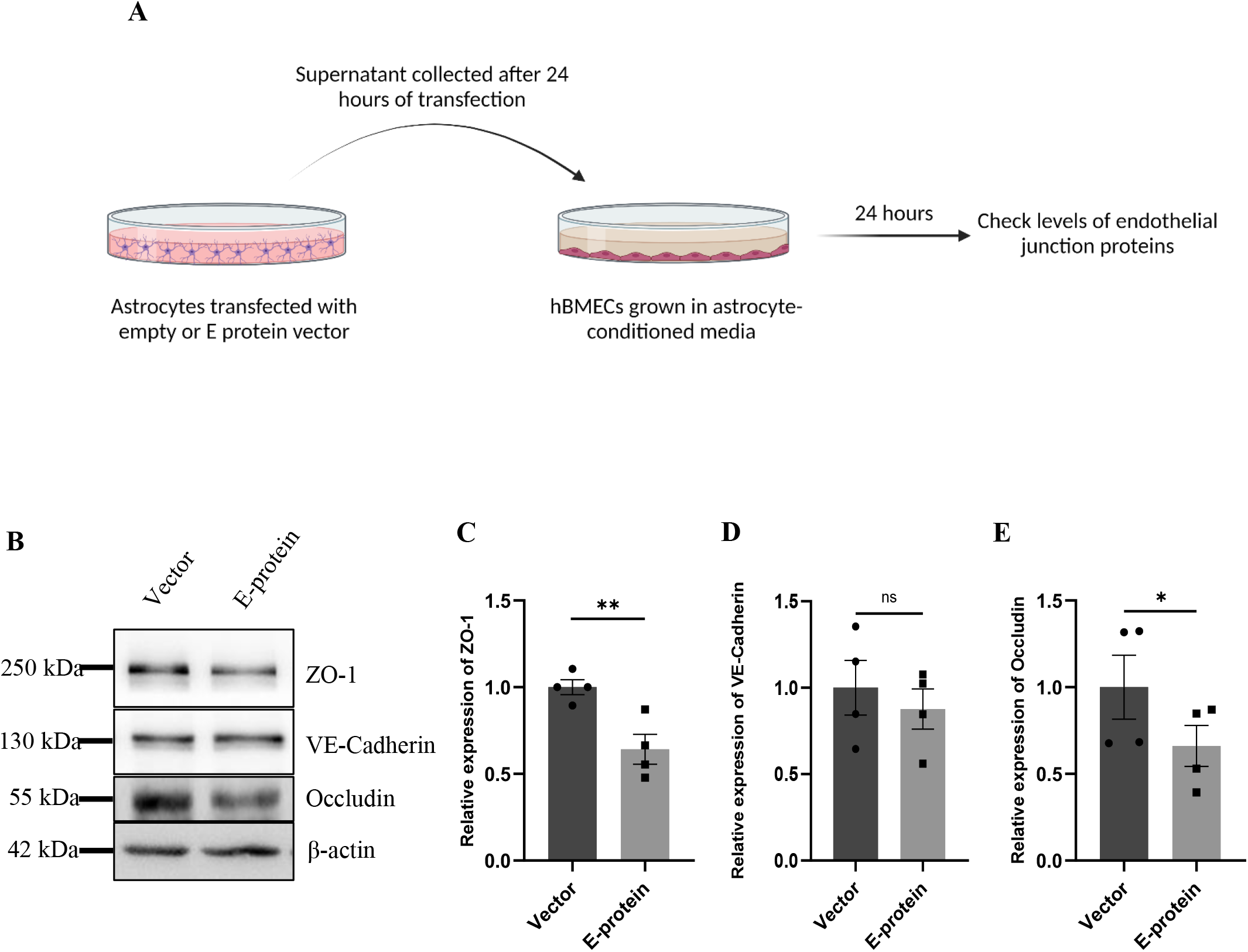
Reactive astrocytes affect endothelial junction proteins in BMECs. Astrocytes were transfected with empty vector and E protein for 24 hours. Schematic representation of experimental paradigm followed (A). Supernatants were collected and mixed with equal amount of fresh endothelial growth media. Brain microvascular endothelial cells were exposed to the astrocyte-conditioned media for 24 hours. (B) Total protein was isolated and subjected to Western blotting against endothelial junction proteins-ZO-1 (C), VE-Cadherin (D), and occludin (E). β-actin was used as loading control. One representative blot is shown. Bar graphs shows relative protein expression in indicated groups. Data represents mean ± SEM for at least three independent experiments. *p<0.05, **p<0.01, ns= not significant, with respect to control.

**Figure 8:**
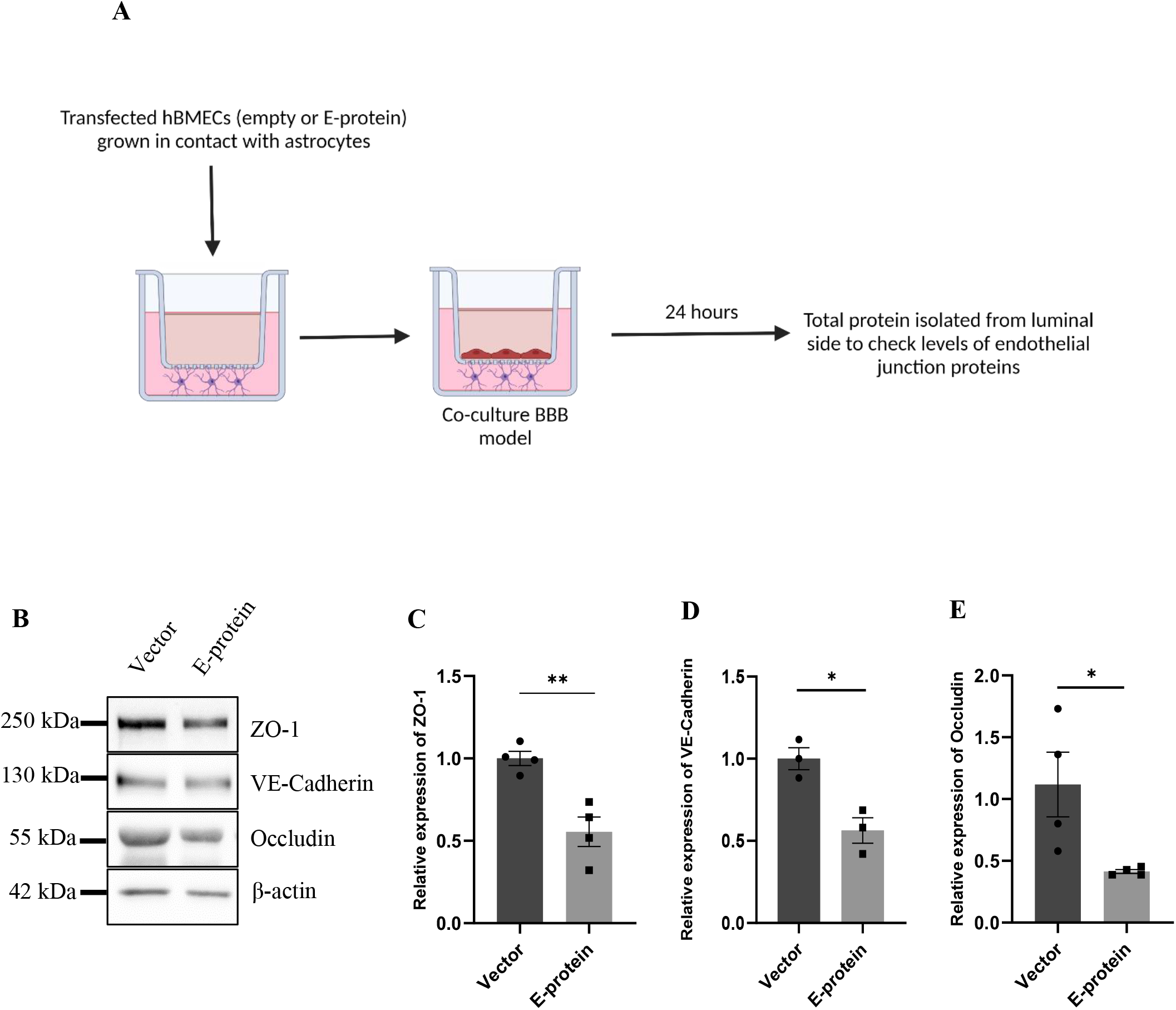
ZIKV E protein expression in BMECs affects endothelial proteins in BBB of astrocytes and BMECs. Transfected BMCEs were grown in contact with astrocytes for 24 hours on transwell apparatus to form co-culture of BBB. Schematic representation of experimental paradigm followed (A). (B)Total protein was isolated from the luminal side of transwell and subjected to Western blotting against endothelial junction proteins-ZO-1 (C), VE-Cadherin (D), and occludin (E). β-actin was used as loading control. One representative blot is shown. Bar graphs shows relative protein expression in indicated groups. Data represents mean ± SEM for at least three independent experiments. *p<0.05, **p<0.01 with respect to control.

## Discussion

Our study aimed at the investigation of role of principal surface protein of Zika virus i.e., envelope (E) protein on the characteristic properties of cells comprising the BBB, and delineate the molecular mechanisms for the same. Majority of work in the field has been mostly done in monoculture BBB model systems, here we employed a two-dimensional model of hBMECs and astrocytes to recapitulate the *in vivo* physiology of BBB. In our study we have employed two *in-vitro* model systems of BBB-1) monolayer of primary hBMECs cultured on transwell and 2) a contact-based co-culture of primary hBMECs and human neural stem cell derived astrocytes cultured on either side of transwell apparatus. Using both of our model systems we found that exposure of E protein to the BBB model resulted in significant modulation of BBB integrity and endothelial permeability. However, the changes in hBMECs permeability changes were small, they were significant to induce cellular activation of hBMECs. We further investigated the molecular drivers of BBB disruption induced by ZIKV E protein. We studied the effect on expression of tight and adherens junction proteins which are responsible for maintaining the junctional stability. Indeed, ZIKV E protein altered expression of endothelial junction proteins - ZO-1, Occludin and VE-Cadherin in both monoculture and co-culture model systems. Previous studies also showed a downregulation of tight junction protein expression in hBMECs infected with ZIKV Asian strain (PRVABC9, Puerto Rico), when compared to African strains (R103451 and MR 766)^32^. Our study further fills the lacunae of other *in vitro* components of BBB model as seen by presence of astrocytes in contact with hBMECs. Our contact-based two-dimensional model system of human origin serves as a more fitting *in vitro* model system to study effect of ZIKV proteins, which has not been reported before.

During viral invasion and BBB disruption BMECs function as source of pro-inflammatory chemokines and cytokines^33^. Hence, we tested the hypothesis if ZIKV E protein has any influence on BMECs activation. Our experiments describe activation of BMECs on E protein exposure, proinflammatory cytokines and chemokines (IL-6, IL-8, IL-1β, CCL2, CCL5, and CXCL10) which are known to alter BBB integrity upon CNS infection are increased^33–37^. Along with increase in inflammatory molecules, BMECs showed increased levels of other endothelial barrier disrupting molecules such as cell adhesion molecules-ICAM-1 and VCAM-1. Increased CAM levels are known to augment viral entry contributing in immune cell infiltration and elevate inflammatory response at the site of BBB^38^. Endothelium permeabilization is one mechanism by which neurotropic viruses access the CNS^39^. Dysregulation of endothelial junction proteins, increased expression of cell adhesion molecules and inflammatory molecules can influence the viral access into the CNS without completely disrupting the BBB^40^. Our findings are in agreement with the previous studies showing ZIKV persistence in hBMECs and potentially use of the paracellular route to enter the privileged neuronal compartments^41^.

Astrocytes are found to be major brain cell population susceptible to ZIKV infection during fetal brain development^42^. Their close contact with BMECs in maintaining the neurovasculature makes them eminent during viral infections^43^. Infected astrocytes are known to show cytopathogenic effects in response to ZIKV infection and act as reservoir for viral replication^44^. We investigated how astrocytes in BBB respond in presence of ZIKV E protein. Astrocytes transfected with E protein were found to be highly reactive as reflected by increase in GFAP and vimentin expression. Astrocytes in their reactive state exhibited increased transcript levels of pro-inflammatory cytokines like IL-6 and IL-8. MCP-1 being an important chemokine in the inflammatory events in viral invasion was found to be increased when astrocytes were exposed to ZIKV E protein. MCP-1 is also known to regulate BBB permeability as brain endothelial cells express MCP-1 receptor, CCR2^45,46^. Glutamate homeostasis was also perturbed resulting in increased levels of glutamate in the extracellular milieu of transfected astrocytes. These present findings indicate that activated astrocytes play crucial role in disruption of BBB integrity during the course of ZIKV infection. Furthermore, our co-culture BBB model system closely mimicked the physiological environment where both the cells were in mutual contact and their individual contributions to the BBB pathophysiology was studied in response to ZIKV E protein.

The attained neurotropic nature of ZIKV makes it capable to infect almost all types of brain cells – microglia, neural progenitor cells and astrocytes; resulting in severe consequences during foetal brain development^19,47^. The associated neurological complications due to vertical transmission (from infected pregnant women to foetus) mainly involves miscarriages, impaired foetal growth or foetuses born with congenital zika syndrome (CZS)^48^. Studies have demonstrated incidence of ZIKV infection post breaching the blood-placental barrier and blood-brain barrier^49^. Clinical studies indicated prevalence of cognitive abnormalities like seizure disorders and motor impairments in children born with CZS^50,51^. As the membrane fusion and subsequent entry of the virus is facilitated by E protein, it proves to be a conducive target for drug therapy and vaccine development^52,53^. Our previous lab studies have focused on how ZIKV E protein perturbed the miRNA circuitry of human fetal neural stem cells (fNSCs) pertaining to cell cycle arrest and inhibition of proliferation^19,20^. This strongly highlights the fact that E protein plays eminent role in pathogenicity of the virus. In addition, endothelial dysfunction has been also reported due to ZIKV NS1 protein bystander effects. NS1 protein has been shown to destabilize VE-cadherin complex and promoted disruption of claudin 5 (CLDN5) affecting the endothelial integrity^54,55^. To further accentuate the findings resulting from single ZIKV structural protein, our study further requires the validation that can be performed in human ZIKV microcephalic brain tissues. Also, even though we tried to closely mimic the physiological environment by establishing a contact -based model of hBMECs and astrocytes, our present study will be benefitted from *in vivo* ZIKV animal models to better understand the corroborated effects of ZIKV E protein. Altogether, our study concludes that ZIKV E protein comprises the BBB maintenance. Our findings implicates that the envelope protein of ZIKV dysregulates the properties of hBMECs which may aid in release of viral particles into the parenchyma without complete disruption of BBB. The altered state of BBB results from the dysregulated state of endothelial junction proteins and activated state of BMECs, resulting in an inflammatory cascade. In addition to this, astrocytes were also found to be susceptible and reactive towards exposure of E protein, thereby influencing the BBB integrity suggesting a potential mechanism in ZIKV-mediated BBB dysregulation through direct contact with hBMECs.

