## Supplementary figures for "Zika virus E protein alters blood-brain barrier by modulating brain microvascular endothelial cell and astrocyte functions"

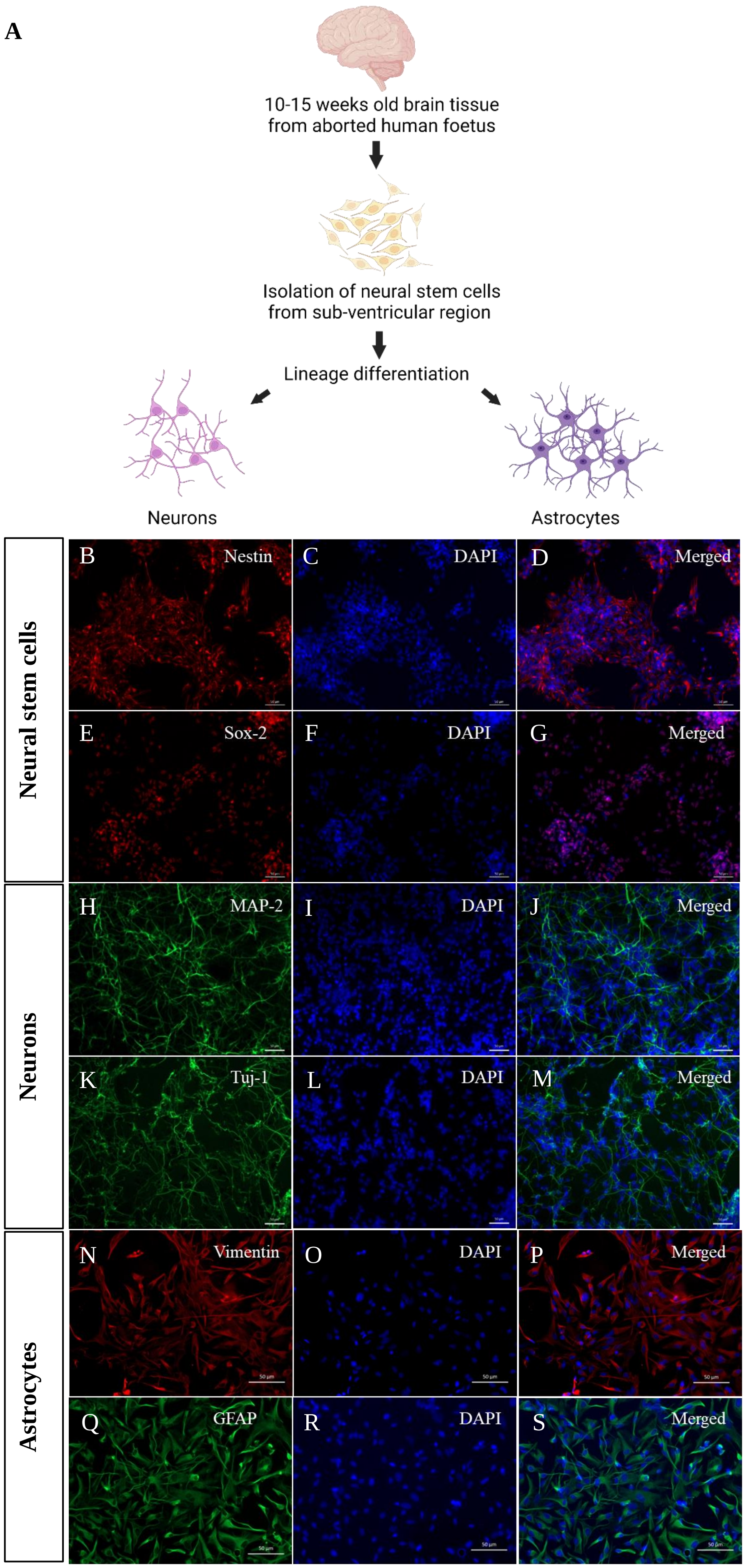


**Supplementary figure. 1 Isolation of neural stem cells (NSCs) and characterization of lineage specific markers:** Schematic representation of isolation of NSCs from aborted human foetus and lineage differentiation (A). (B-S) Immunocytochemistry images of functional markers for neural stem cells (Nestin, Sox-2), neurons (MAP-2, Tuj-1) and astrocytes (Vimentin, GFAP) in indicated groups. DAPI (blue) for nucleus. Scale bar 50 µm.

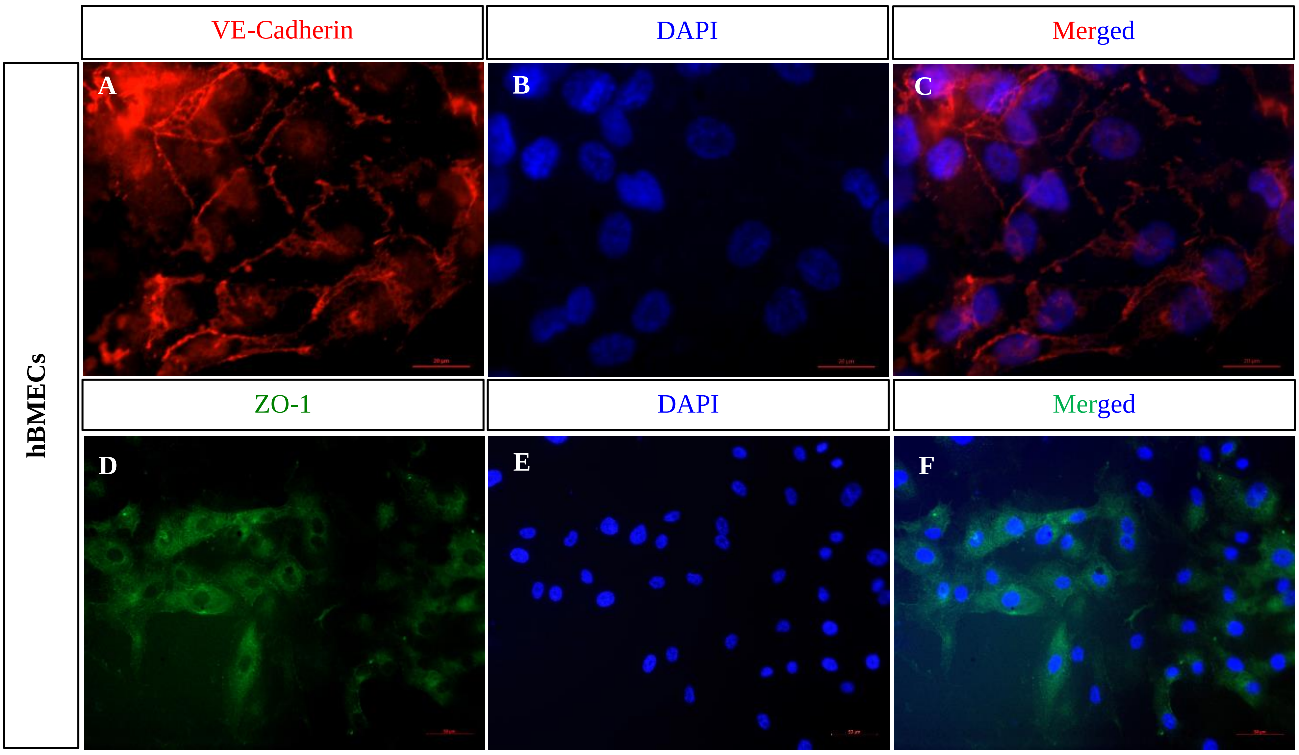


**Supplementary figure. 2 Human primary brain microvascular endothelial cells (BMECs) as *in vitro* model system.** hBMECs were immunostained with endothelial specific markers- VE-Cadherin (A-C, red), ZO-1 (D-F, green), and DAPI (blue) for nucleus. Scale bar = 50 µm.

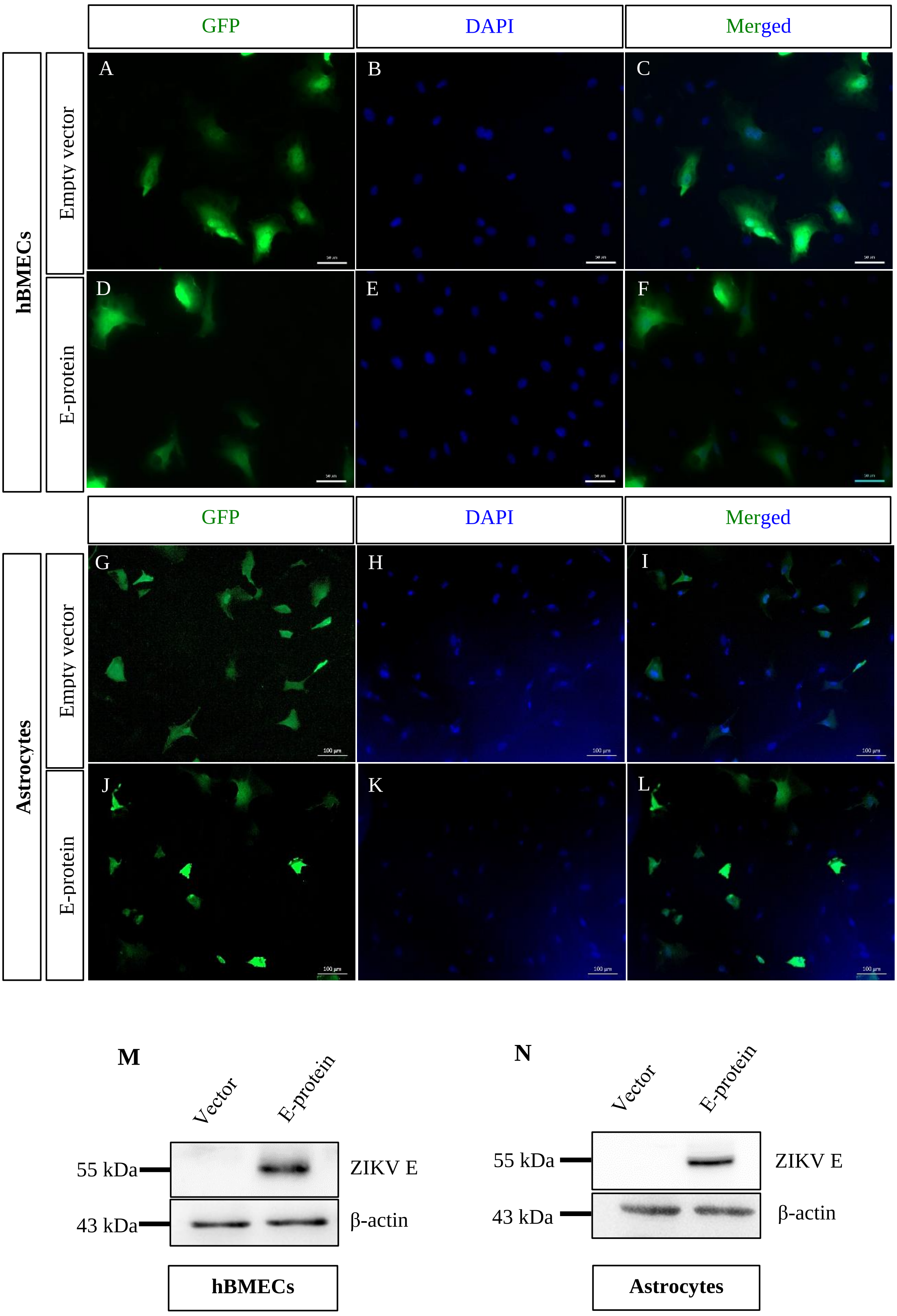


**Supplementary figure. 3 Transfection efficiency of plasmids:** (A-F) Immunocytochemistry images of BMECs showing GFP-positive cells (green) and DAPI (blue) for nucleus in indicated groups (Empty vector and E-protein) post 24-hour transfection. Scale bar 50 µm. (G-L) Immunocytochemistry images of astrocytes showing GFP-positive cells (green) and DAPI (blue) for nucleus in indicated groups (Empty vector and E-protein) post 24-hour transfection. Scale bar 100 µm. (M-N) Western blotting against ZIKV E-protein showing expression of E-protein in indicated groups post 24-hour transfection in BMECs and astrocytes, respectively. β-actin was used as a loading control.
