## Supplementary table for "Zika virus E protein alters blood-brain barrier by modulating brain microvascular endothelial cell and astrocyte functions"

| | |
|---|---|
| IL-6 FP | 5’-GGAGACTTGCCTGGTGAAA -3’ |
| IL-6 RP | 5’-CTGGCTTGTTCCTCACTACTC -3’ |
| IL-8 FP | 5’-GGACAAGAGCCAGGAAGAAA-3’ |
| IL-8 FP | 5’-ACACAGAGCTGCAGAAATCA-3’ |
| IL-1β FP | 5’-CAAAGGCGGCCAGGATATAA-3’ |
| IL-1β RP | 5’-CTAGGGATTGAGTCCACATTCAG-3’ |
| CCL2 FP | 5’-GGCTGAGACTAACCCAGAAAC-3’ |
| CCL2 RP | 5’-GAATGAAGGTGGCTGCTATGA-3’ |
| CCL5 FP | 5’-CTCCGTCACAACAACAACAAC-3’ |
| CCL5 RP | 5’-AGAGCTCAGAACCTAGAGACTT-3’ |
| CXCL10 FP | 5’-ACCAAATCAGCTGCTACTACTC-3’ |
| CXCL10 RP | 5’-CAGGGTCAGAACATCCACTAAG-3’ |
| ICAM-1 FP | 5’-CCGCAGTCATAATGGGCACT-3’ |
| ICAM-1 RP | 5’- GGTTTCATGGGGGTCCCTTT-3’ |
| VCAM-1 FP | 5’-GGGAAGCCGATCACAGTCAA-3’ |
| VCAM-1 RP | 5’- TCCTGTCTGCATCCTCCAGA-3’ |
| PTGS-2 FP | 5’- TGTATGAGTGTGGGATTTGACC -3’ |
| PTGS-2 RP | 5’-TGTGTTTGGAGTGGGTTTCAG -3’ |
| GFAP FP | 5’-ACCTGCAGATTCGAGAAACCAG-3’ |
| GFAP RP | 5’-TAATGACCTCTCCATCCCGCATC-3’ |
| Vimentin FP | 5'- AAGTCCGCACATTCGAGCAA-3' |
| Vimentin RP | 5'-CTACCAACTTACAGCTGGGC-3' |
| GAPDH FP | 5’-CAAGAGCACAAGAGGAAGAGAG -3’ |
| GAPDH RP | 5’-CTACATGGCAACTGTGAGGAG -3’ |

**Supplementary table 1:**

List of qPCR primers used in the study.
